# Underlying environmental factors and geomagnetic fields linked to 45 pilot whales *Globicephala macrorhynchus* stranding in Modung white beach, Indonesian coast on 19^th^ February 2021

**DOI:** 10.1101/2021.02.25.432840

**Authors:** Andri Wibowo

## Abstract

The reason whale and dolphin stranding is not fully understood and it is not linked to a standalone variable. Theories assume intertwined factors including sickness, underwater noise, navigational error, geographical features, the presence of predators, poisoning from pollution or algal blooms, geomagnetic field, and extreme weather are responsible to whale stranding. On 19^th^ February 2021, a pod consists of 45 pilot whales *Globicephala macrorhynchus* was stranded in a remote 7050 m^2^ Modung white beach of Indonesian coast. This paper aims to assess the environmental factors that may be can explain and link to this stranding cases. Those factors include bathymetry, plankton cell density measured using MODIS, water sediment load measured using Sentinel 2 Bands 4,3,1, vessel traffic, precipitation (inch) and thunder (CAPE index J/kg), water salinity and temperature, and geomagnetic field (nT). The results show the water near stranding sites were shallow, has sediment load, high plankton density, warmer, receiving torrential rain prior stranding, having weak geomagnetic field and high total magnetic field change/year. The combination of those environmental covariates may influence the behavior, navigation, and echolocation of the said stranded pilot whale.

## Introduction

Cetacean stranding cases are growing recently. This phenomena is an intertwined of multiple factors involving complex combinations of biological, physiological, and environmental factors. Several studies have confirmed the stranding cases were related to biological and physiological of a whale. A study has noticed that stranding cases may be related to the fatigue. It is assumed that there was large amounts of energy that dolphins and whales expend when fleeing threats may be linked to mass strandings. Whale used more energy to swim fast than to cruise at normal speeds when avoiding threats caused by people as well as predators. This escape requires energy (Williams et al 2017) and could come at a cost to mammals that live in the seas. It is probably the whales escape the predator and stranded in the coast. Since escaping from predator require energy, the whales do not have adequate energey to swim back from beach to deep sea.

Regarding biological factors, stranded whales were also related to the age and even disease. Old whales may find it difficult to keep up with their pod or resist heavy swells or inshore currents. Because of failing strength these animals may strand. Whales with poor health conditions in spite of disease and malnutrition are also can experience stranding. Poor healths are related to the malnutrition and food shortage due to overfishing, ingestion of plastic, and consmuptions of natural toxins due to harmful algal blooms. The whales with poor health can be picked up by a wave and thrown onto a beach.

Indonesia is a country with a vast coastline. This 95181 km coast line recenty has been a site of whale stranding that happens frequently. Then this paper aims to assess the particular environmental factos that related to the whale stranding cases recently https://abcnews.go.com/International/wireStory/45-pilot-whales-survive-mass-stranding-indonesia

## Materials and Methods

### Study area

The study area is a remote coastal area located in south of Madura island. This island is located in the north of Java island. On 19^th^ February 2021 at 16.00 pm, 45 pilot whales *Globicephala macrorhynchus* were reported stranded in this coast. The exact location was a fragment of white beach sizing 7050 m^2^ surrounded by road, vegetation, and brackish fish pond.

### Environmental factors

Several environmental factors were collected. The collection time was 3 hours before the stranding from 13.00 pm to 16.00 pm on 19^th^ February 2021. The environmental factor data were retrieved through the remote sensing and environmental data providers. The retrieved environmental data including bathymetry, plankton cell density using MODIS, water sediment load using Sentinel 2 Bands 4,3,1, vessel traffic, precipitation (inch) and thunder (CAPE index J/kg), water salinity and temperature, and geomagnetic field (nT).

## Results and Discussion

### Coast bathymetry

Figure 2 presents the bathymetry profile of Modung beach. It can be seen that the water around the beach was dominated by shallow water with depth ranging from 0 to 9 m. For whales, bathymetry is one of determinant factors influencing the whale distribution and habitat preferences. Bathymetric features representing continental and reef slopes, shallow banks and seamounts tend to be areas contain high marine productivity, in particular high zooplankton abundance, and this lead to the predator prey aggregations including whale (Afonso et al. 2014). Bouchet et al. (2015) and Copping et al. (2018) observed areas with complex bathymetry including seamounts or steep slopes found on outer reefs accumulate zooplankton that subsequently attracts filter feeders inhabiting epipelagic and mesopelagic depths. This was confirmed by Sims (2008) with correlations of basking sharks abundance, steeper slopes, and zooplankton density.

**Figure 1.**
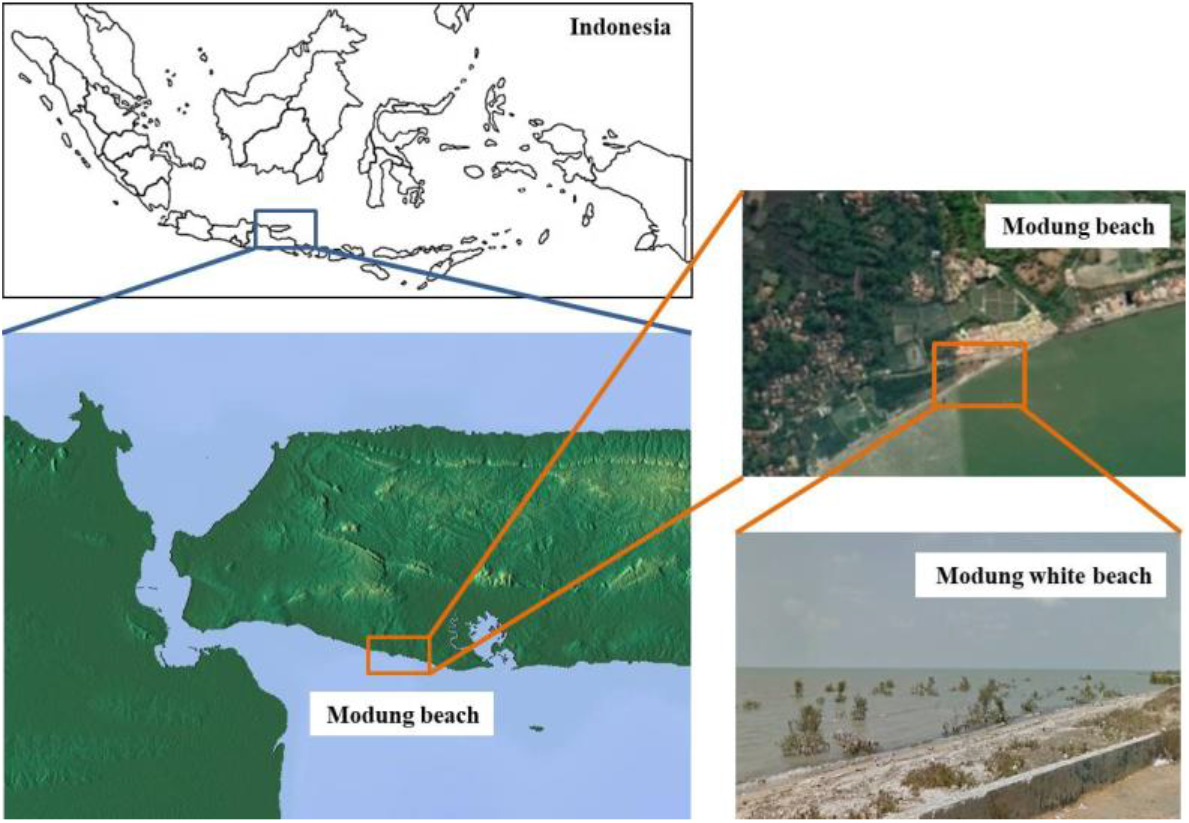
Location of 45 pilot whales *Globicephala macrorhynchus* stranding in 7050 m^2^ Modung white beach, south of Madura island on 19^th^ February 2021 at 16.00 pm

**Figure 2.**
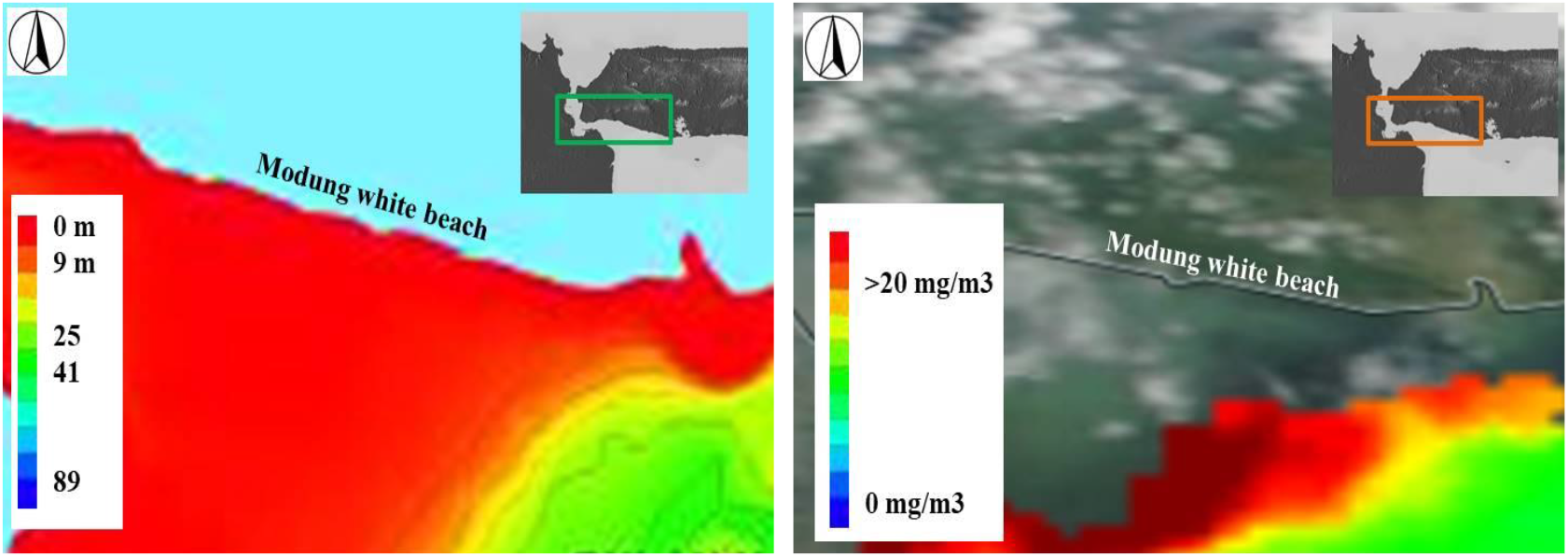
Bathymetry (left) and plankton density (mg/m^3^) measured using MODIS and distribution (right) near Modung beach

For whale shark, coastal areas of shallow bathymetry in close proximity to the reef slope and deeper water were preferred habitats and whales were common found in this area (De la Parra Venegas et al. 2011, Donati et al. 2016, Diamant et al. 2016). This whale behavior is in accordance with the stranding case of pilot whale in this study where the whale aggregations were found in shallow water.

### Plankton abundance and HAB potential

The MODIS imagery shows the plankton abundance in offshore of Modung beach (Figure 2). The plankton density was observed at its maximum values with 20 mg/m^3^. This high density shows the water near the Modung is rich and a potential feeding grounds including for top predator like pilot whale. Short-finned pilot whales feed on squids and fishes that in turn prey on micronektonic organisms. The stranding event on 19^th^ February 2021 was preceded by an aggregation of pilot whale driven by the presences of planktonic organism at high density near shore. This is comparable to Abécassis et al (2015) findings where whales and micronekton were found more inshore. Whereas the high density of plankton is not always a benefit for whales considering the potential of harmful algal bloom (HAB).

Plankton density and related HAB can be a variables linked to the whale stranding. HAB can pose a threat to marine mammal including whale. Beyond human causes, mass whale stranding have been attributed to herding behaviour, large-scale oceanographic fronts, and HABs (Pyenson et al 2014). Because algal toxins cause organ failure in marine mammals, HAB is one of the most common whale stranding underlying factors with broad geographical and widespread taxonomic impact. HAB enters whale body indirectly through whale preys. It was estimated that a whale consuming 4% of its body weight to prey on mackerel daily that would have contained saxitoxin (STX). This toxin produced by dinoflagellate *Alexandrium*, *Gymnodinium*, and *Pyrodinium* may explain mass-strand of 14 humpback whales (*Megaptera novaeangliae*) within 5 weeks with a stomachs full of undigested Atlantic mackerel (*Scomber scombrus*) (Fire & Van Dolah 2012). Evidence of the whales’ exposure to HAB toxins can be observed in trace levels of paralytic shellfish toxins (PSTs) and domoic acid (DA) in tissues of some dead whales, and fragments of *Pseudo-nitzschia* spp. frustules in whale feces (Wilson et al. 2015). Humpback whales (*Megaptera novaeangliae*) were also reported positive at the highest rates for STX and and domoic acid (DA) (Fire et al 2021).

### Sediment and navigational error

The water near the Modung beach was characterized by sediment loads (Figure 3). The water receives the sediment loads from the river located in the west in Java island. The river discharges suspended materials from the Java mainland in the west. Whales and dolphins are animals that depend on the echolocation to navigate in the water. Whereas shallow and sloping shores consisted of soft sediment can confuse the echolocation used by whales and dolphins to find their way around. In Modung beach it is estimated that the pilot whales foraged in shallow coast to follow the plankton density and their echolocation was disrupted due the presences of massive sediment loads in water discharged from the mainland. Sediment loads in water can increase turbidity that exceeds natural level. Sediment can also contain pollutant that can harm whales (Victoria. et al 2015). Nearshore sediment loads are related to the anthropogenic activities in the upstream areas. Massive settlement developments, deforestation, and poor sediment management in the river mouth will accelerate the sediment load in offshore.

**Figure 3.**
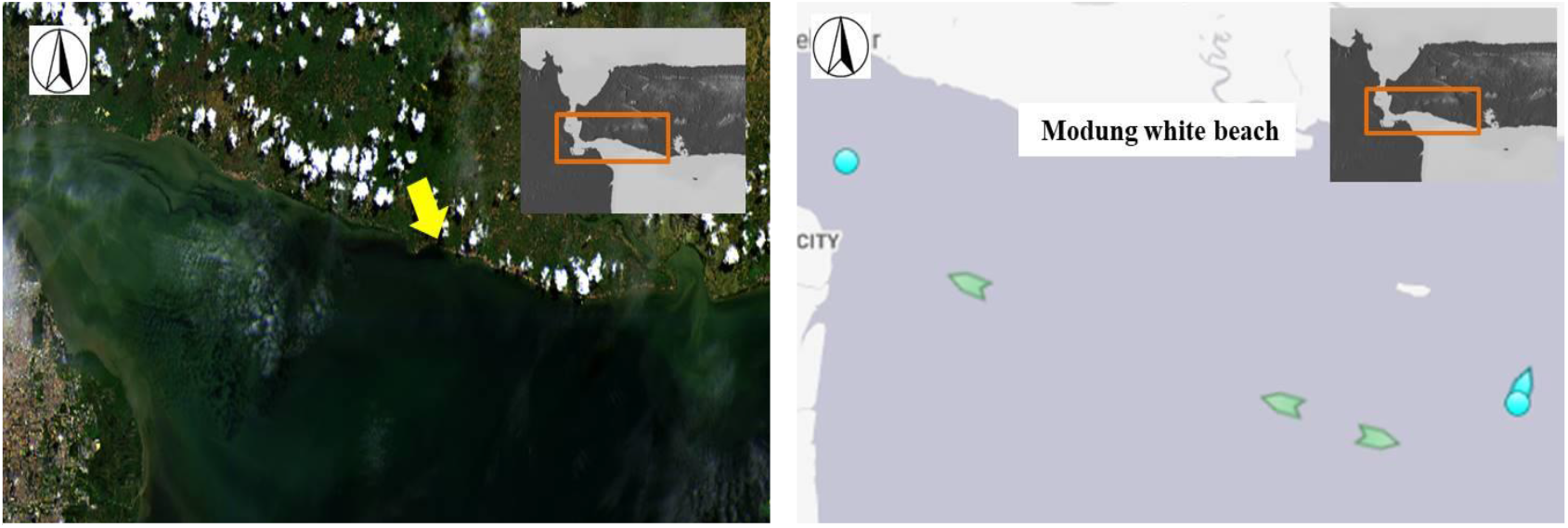
Sediment load in water (left) analyzed using Sentinel 2 Bands 4,3,1 and vessel traffics (right) near Modung beach (yellow arrow)

### Vessel traffic and noise

Another anthropogenic cause that can contribute to the whale stranding is the vessel traffic. The presence of vessel traffic can contribute to the direct impact including collision and even disrupting the whale echolocation. The Figure 3 presents the vessel traffic condition near Modung beach. It is clear that the traffic is not that busy. Whereas the water is used as shipping routes for large vessel to go from and to the ports in the Java mainland. Neilson et al (2012) have reported 108 whale-vessel collisions in for period of 1978–2011 in Alaska. From that cases, 25 cases are known to have resulted in the whale’s death. Most collisions involved humpback whales (86%) with six other species documented. Common collision (60%), involves small vessel sizing <15 m, medium (15–79 m, 27%) and large (≥80 m, 13%) vessels also struck whales. Even in some areas, one-third of all fin whale and right whale strandings appear to involve ship strikes as reported by Laist et al (2001).

Besides causing direct impact on whale including collisions, vessel traffic with its noise can also contribute to the whale echolocation disruptions. The number of vessel is on the rise ranging from private boats in coastal areas to commercial ships crossing oceans. This rise has led to a concomitant increase in underwater noise has been reported in several regions around the globe (Christine et al 2019). The effects of underwater noise from vessel traffic on whale include behavioral responses, acoustic interference (i.e., masking), temporary or permanent shifts in hearing threshold (TTS, PTS), and stress. Jensen et al (2009) using experiments have confirmed that pilot whales in a quieter deep-water habitat could suffer a reduction in their communication range of 58% caused by a vessel at similar range and speed.

### Precipitation and thunder noises

Besides noise generated from vessel traffics, underwater noise is also generated by precipitation and thunderstorm. Natural noises produced by sea waves breaking on the coast, currents moving over reef, raindrops on the ocean surface, tides, oceanic turbulence and the sound produced by seaquakes and submarine volcano eruptions are typical natural background sounds that also can impact marine organisms (Peng et l. 2015). Before whale stranding happens at 16.00 pm, Modung beach was experiencing torrential rain with thunderstorm (Figure 4). When the rain presences and the number of heavy rain fall drops increases, it can dominate the noise levels that are produced by the rain drops. The acoustic noise level of large and very large rain drops can produce an approximately straight fit line in the frequency spectrum range less than 5 kHz. The frequency spectra that were generated by the rain noise were in the range of 2-7 kHz. The frequency spectrum is mainly produced by the rain drops of size in the range between 2.2 mm to 4.6 mm in diameter. The result measured by Ashokan et al (2015) shows that the noise spectrum level increases with precipitation and the correlation between the noise level and precipitation is very good in the frequency of 2-7 kHz.

**Figure 4.**
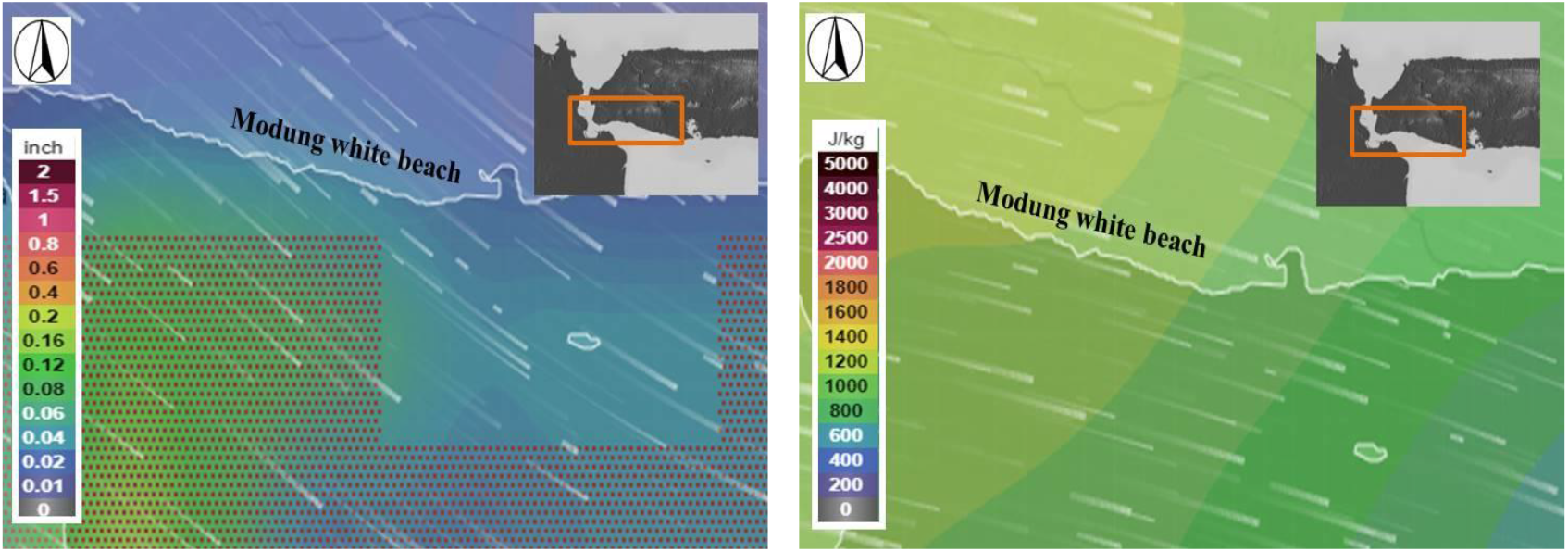
Precipitation (left) and thunderstorm (CAPE index J/kg) intensities (right) prior whale stranding in water near Modung beach

### Water temperature and salinity

Figure 5 presents the water temperature and salinity near Modung beach water prior to whale stranding. It can be seen that the water was warmer and 2°C higher than surrounding water. The water near whale stranding sites was having salinity very low and closed to the freshwater. The warmest ocean temperatures can contribute to the whale stranding behavior. A spike in temperatures is affecting where the prey is moving and as a consequence prey moving and whale species following. Warming sea temperatures lead to multiple changes in the seascape including a major shift in the biological, chemical, and physical characteristics of the habitats that whales inhabit would eventually affect whales prey availability and physiology

**Figure 5.**
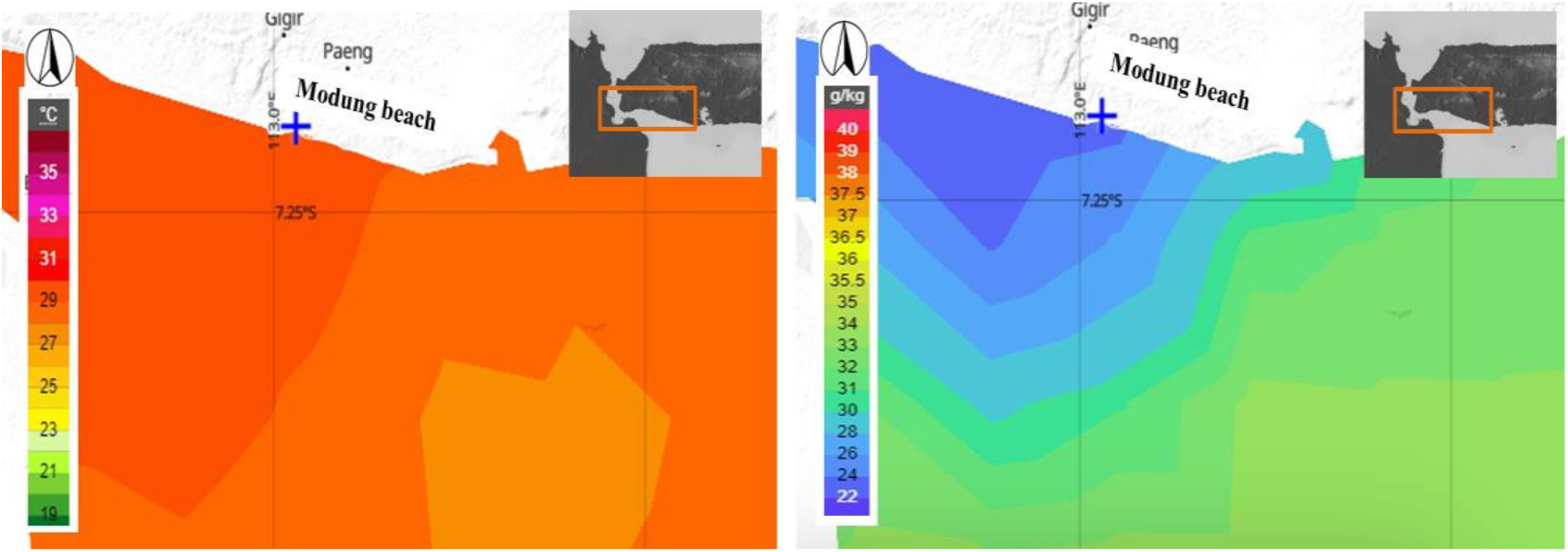
Water temperature (°C) (left) and salinity (g/kg) values (right) prior whale stranding in water near Modung beach

### Geomagnetic field

Figure 6 presents the geomagnetic value occurred in water near Modung beach. Whales have magnetic sensory and appear to use the flux density of the earth’s magnetic field (total field) in two ways as an aid to travel (Klinowska et al 1990). The topography of the local magnetic field is used as a map, with the whales generally moving parallel to the contours. Geomagnetic field can provide a great deal of stable positional and directional information (Horton et al. 2020).

**Figure 6.**
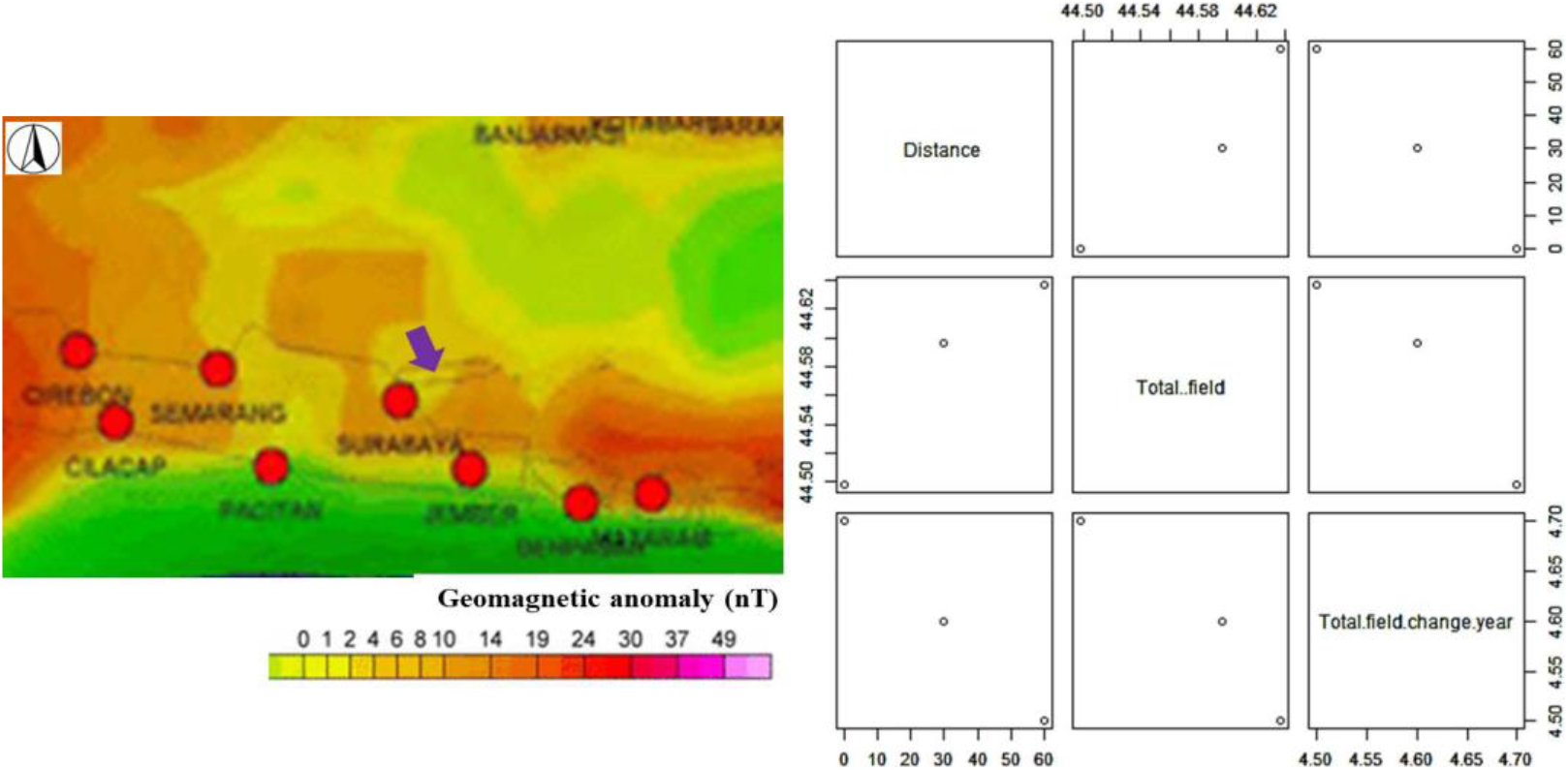
Geomagnetic anomaly (nT) (left) and correlation of distance to water (km), total field (nT) and total field change/year (nT/yr) measured on 19^th^ February 2021 (right) in water near Modung beach

A weak and low magnetic field will cause the whale becomes disoriented and stranded in the beach. Walker et al. (1992) observed that whale routes were correlated with maximum values of geomagnetic fields. While Kirschvink et al. (1986) found that local magnetic minima was correlated with highly significant tendencies for whales to beach themselves near coastal locations. In Modung, the geomagnetic field inshore near the beach with a distance to water was 0 km was lower than offshore. Whereas the annual flux was higher inshore than offshore. It indicates that the geomagnetic fields are weaker near the beach and cause the whale loss its disorientation. Recent study have found that intertwined effects of geomagnetic and solar storm may impact the whales become disoriented and lead to mass stranding (Granger et al 2020).

## Conclusion

The reason whale and dolphin stranding is not fully understood and it is not linked to a standalone variable. The results from this study show the water near stranding sites in Modung beach were shallow, has sediment load, high plankton density, warmer, receiving torrential rain prior stranding, having weak geomagnetic and high total magnetic field change/year. The combination of those environmental covariates may influence the behavior, navigation, and echolocation of the said stranded 45 pilot whales happens on the last 19^th^ February 2021.

## Notes

### Competing Interest Statement

The authors have declared no competing interest.

## References

Abécassis M, Polovina J, Baird R, Copeland A, Drazen J, Domokos R, Oleson E, Jia Y, Schorr G, Webster D, Andrews R. 2015. Characterizing a Foraging Hotspot for Short-Finned Pilot Whales and Blainville’s Beaked Whales Located off the West Side of Hawai’i Island by Using Tagging and Oceanographic Data. PloS one. 10. e0142628. 10.1371/journal.pone.0142628.

Afonso P, McGinty N, Machete M. 2014. Dynamics of whale shark occurrence at their fringe oceanic habitat. PLOS ONE 9(7).

Andrady AL 2015 Persistence of Plastic Litter in the Oceans. In: Bergmann M., Gutow L., Klages M. (eds) Marine Anthropogenic Litter. Springer, Cham.

Ashokan M, Latha G, Ramesh R. 2015. Analysis of shallow water ambient noise due to rain and derivation of rain parameters Applied Acoustics 88: 114–122

Bouchet PJ, Meeuwig JJ, Kent S, Chandra P, Letessier TB, Jenner CK. 2015. Topographic determinants of mobile vertebrate predator hotspots: current knowledge and future directions. Biological Reviews 90(3):699–728

Christine E, Sarah M, Schoeman RP, Smith JN, Trigg LE, Embling CB. 2019 The Effects of Ship Noise on Marine Mammals—A Review Frontiers in Marine Science 6

Copping JP, Stewart BD, McClean CJ, Hancock J, Rees R. 2018. Does bathymetry drive coastal whale shark (Rhincodon typus) aggregations? PeerJ 6:e4904

De la Parra Venegas R, Hueter R, Cano JG, Tyminski J, Remolina JG, Maslanka M, Ormos A, Weigt L, Carlson B, Dove A. 2011. An unprecedented aggregation of whale sharks, Rhincodon typus, in Mexican coastal waters of the Caribbean Sea. PLOS ONE 6(4):e18994–e18994

Diamant S, Pierce SJ, Ramírez-Macías D, Heithaus MR, D’Echon AG, D’Echon TG, Kiszka JJ. 2016. Preliminary observations on whale sharks in Nosy Be, Madagascar. In: QScience Proceedings. 15

Donati G, Rees RG, Hancock JW, Jenkins TK, Shameel I, Hindle K, Zareer I, Childs A, Cagua EF. 2016. New insights into the South Ari atoll whale shark, Rhincodon typus, aggregation. In: QScience Proceedings. 16

Fire SE, Van Dolah FM. 2012 Marine Biotoxins: Emergence of Harmful Algal Blooms as Health Threats to Marine Wildlife in ew Directions in Conservation Medicine: Applied Cases in Ecological Health, editedby A. Alonso Aguirre, Richard S. Ostfield, and Peter Daszak

Fire SE, Bogomolni A, DiGiovanni RA Jr, Early G, Leighfield TA, Matassa K, et al. 2021 An assessment of temporal, spatial and taxonomic trends in harmful algal toxin exposure in stranded marine mammals from the U.S. New England coast. PLoS ONE 16(1): e0243570

Granger J, Walkowicz L, Fitak R, Johnsen S. 2020. Gray whales strand more often on days with increased levels of atmospheric radio-frequency noise Current Biology 30

Horton TW, Zerbini AN, Andriolo A, Danilewicz D and Sucunza F 2020 Multi-Decadal Humpback Whale Migratory Route Fidelity Despite Oceanographic and Geomagnetic Change. Front. Mar. Sci. 7:414.

Jensen F, Bejder L,Wahlberg M, Aguilar de Soto N, Madsen P 2009. Vessel noise effects on delphinid communication. Marine Ecology Progress Series, 395: 161–175.

Kirschvink JL, Dizon AE, Westphal JA. 1986. Evidence from Strandings for Geomagnetic Sensitivity in Cetaceans Journal of Experimental Biology 120: 1–24

Klinowska M. 1990 Geomagnetic Orientation in Cetaceans: Behavioural Evidence. In: Thomas J.A., Kastelein R.A. (eds) Sensory Abilities of Cetaceans. NATO ASI Series (Series A: Life Sciences), vol 196. Springer, Boston, MA.

Laist D, Knowlton A, Mead J, Avenue C, Collet A, Podestà M. 2001. Collisions between ships and whales. Marine Mammal Science. Marine Mammal Science. 35–75. 10.1111/j.1748-7692.2001.tb00980.x.

Neilson JL, Gabriele CM, Jensen A, Jackson K, Straley JM. 2012 Summary of Reported Whale-Vessel Collisions in Alaskan Waters Journal of Marine Sciences

Peng C, Zhao X, Liu G. 2015. Noise in the Sea and Its Impacts on Marine Organisms. International journal of environmental research and public health, 12(10), 12304–12323.

Pyenson ND, Gutstein CS, Parham JF, Le Roux JP et al. 2014. Repeated mass strandings of Miocene marine mammals from Atacama Region of Chile point to sudden death at sea. Proceedings. Biological sciences, 281(1781)

Rodhouse PG, Nigmatullin CM. 1996. Role as consumers. Philos Trans R Soc B Biol Sci.:351: 1003–1022.

Sims DW. 2008. Sieving a living: a review of the biology, ecology and conservation status of the plankton-feeding basking shark cetorhinus maximus. Advances in Marine Biology 54:171–220

Walker MM, Kirschvink JL, Ahmed G, Diction. 1992 Evidence That Fin Whales Respond To The Geomagnetic Field During Migration J. exp. Biol. 171: 67–78

Williams TM, Kendall T, Richter BP, Ribeiro-French CR, John JS, Odell KL, Losch BA, Feuerbach DA, Stamper MA 2017 Swimming and diving energetics in dolphins: a stroke-by-stroke analysis for predicting the cost of flight responses in wild odontocetes Journal of Experimental Biology220: 1135–1145

Victoria LG, Todd IB, Todd JC, Gardiner ECN, Morrin NA et al 2015. A review of impacts of marine dredging activities on marine mammals, ICES Journal of Marine Science 72(2): 328–340

Wilson C, Sastre V, Hoffmeyer M, Rowntree V, Fire Sp et al. 2015. Southern right whale (*Eubalaena australis*) calf mortality at Península Valdés, Argentina: Are harmful algal blooms to blame?. Marine Mammal Science. 32. 10.1111/mms.12263.

